# Selective activation of intracellular β1AR using a spatially restricted antagonist

**DOI:** 10.1101/2023.11.22.568314

**Authors:** Federica Liccardo, Johannes Morstein, Ting-Yu Lin, Julius Pampel, Kevan M. Shokat, Roshanak Irannejad

**Affiliations:** Cardiovascular Research Institute, University of California, San Francisco, USA; Department of Cellular and Molecular Pharmacology, University of California San Francisco, San Francisco, USA; Howard Hughes Medical Institute, University of California, San Francisco, USA; Department of Biochemistry & Biophysics, University of California, San Francisco, USA

## Abstract

G-protein-coupled receptors (GPCRs) regulate several physiological and pathological processes and represent the target of approximately 30% of FDA-approved drugs. GPCR-mediated signaling was thought to occur exclusively at the plasma membrane. However, recent studies have unveiled their presence and function at subcellular membrane compartments. There is a growing interest in studying compartmentalized signaling of GPCRs. This requires development of novel tools to separate GPCRs signaling at the plasma membrane from the ones initiated at intracellular compartments. We took advantage of the structural and pharmacological information available for β1-adrenergic receptor (β1AR), an exemplary GPCR that functions at subcellular compartments, and rationally designed spatially restricted antagonists. We generated a cell impermeable β1AR antagonist by conjugating a suitable pharmacophore to a sulfonate-containing fluorophore. This cell-impermeable antagonist only inhibited β1AR on the plasma membrane. In contrast, a cell permeable β1AR agonist containing a non-sulfonated fluorophore, efficiently inhibited both the plasma membrane and Golgi pools of β1ARs. Furthermore, the cell impermeable antagonist selectively inhibited the phosphorylation of downstream effectors of PKA proximal to the plasma membrane in adult cardiomyocytes while β1AR intracellular pool remained active. Our tools offer promising avenues for investigating compartmentalized β1AR signaling in various context, potentially advancing our understanding of β1AR-mediated cellular responses in health and disease. They also offer a general strategy to study compartmentalized signaling for other GPCRs in various biological systems.

## INTRODUCTION

G protein-coupled receptors (GPCRs) are seven-transmembrane receptors that initiate G protein-mediated signal transduction upon ligand binding ^1^. Studies over the past decade have challenged the textbook model of GPCR signaling by revealing that GPCR signaling is not restricted to the plasma membrane but also occurs from subcellular compartments, such as the endosomes, the Golgi apparatus and the nuclear membranes ^2–12^. The distinct role of subcellularly activated GPCR/G protein complexes remains to be fully understood. There are only a handful of studies demonstrating the cellular and physiological significance of GPCR signaling from subcellular locations^12^. This limitation is partly due to lack of tools that would allow separating receptors’ function at each location. Traditional tools for studying endosomal signaling are based on genetic or pharmacological manipulations of machineries such as clathrin and dynamin that regulate receptor trafficking to endosomes^2,7,8,13–15^. These types of manipulations disrupt trafficking of many other receptors and channels; thus, they can provoke unintended consequences or trigger compensatory mechanisms^16–18^. Endosomal delivery of GPCR ligands, using pH-sensitive nanoparticle-encapsulated ligands, was further developed to better determine the consequence of GPCR activation at the endosomes^16,19,20^. Further studies are necessary to improve therapeutic efficacy of nanoparticle-encapsulated ligands^16^. They are also limited to studying endosomal signaling and cannot be applied to functions of GPCRs that are located at the perinuclear/Golgi membranes.

To better pinpoint the consequence of compartmentalized GPCR signaling, our lab has previously taken advantage of nanobodies that were originally utilized for structural studies of monoamine GPCRs such as beta-adrenergic receptors (βARs)^21,22^. This is because these specific nanobodies bind to activated monoamine receptors (i.e. βARs) with high affinity and to the same region as G proteins^6,21,22^. Thus, when targeted to specific compartment they can disrupt receptor coupling to the endogenous G protein through steric occlusion^6,9,11^. Combining these tools with an inducible system, the rapamycin dimerization system, we were able to test the significance of β1AR and D1DR signaling from the plasma membrane and the Golgi^9,11,23^. While these strategies allow precise manipulation of compartmentalized signaling, it has certain shortcomings including lack of selective nanobodies for most GPCRs as well as insufficient delivery for *in vivo* studies. More recently, a similar approach has been developed based on the RGS domain of GRK2, to disrupt GPCR-Gq protein mediated signaling at various subcellular compartments^24^. While these strategies are useful for cell-based studies, they lack specificity for *in vivo* studies.

Given the growing interest in elucidating the physiological consequences of subcellular GPCRs’ activity, there is an unmet need for developing pharmacological agonists and antagonist that could help resolve their spatial function *in vitro* and *in vivo* ^25^. Although, several studies have already used available GPCRs ligands that are relatively hydrophilic and thus predicted to be poorly membrane permeable ^6,9,11,23,26–29^, a number of reports suggest that some of these hydrophilic drugs can be transported across the lipid membrane by cationic or anionic membrane transporters^30–32^. Therefore, the tissue expression pattern of membrane transporters could impact the consequences of subcellular GPCR activities.

In this study, we report the generation of a specially restricted GPCR antagonist, by focusing on β1AR, an exemplary GPCR. We modeled the common structural and chemical features of known β1AR antagonists and conjugated the pharmacologically active part of the antagonist to either sulfonated or non-sulfonated fluorophore, via click chemistry, to generate cell impermeable and permeable versions of the antagonists, respectively. Using a nanobody-based biosensor that allows for the detection of β1AR activity in intact cells^6,9^, we first confirmed the selectivity of these antagonists at inhibiting distinct pool of β1ARs in cardiomyocytes. We then showed that the cell impermeable β1AR antagonist specifically inhibits downstream effectors of PKA that are in the vicinity of the plasma membrane, without affecting PKA effectors that are known to be regulated by Golgi-localized β1AR signaling^33^. This contrasts with a cell permeable antagonist that inhibits both receptor pools. These drugs can be potentially employed in several biological systems and could allow the interrogation of compartmentalized β1AR signaling for *in vitro* and *in vivo*.

## RESULTS

### OATP1A2 facilitates transmembrane transport of sotalol in murine cardiomyocytes

We have previously shown that β1AR can become activated and couple to Gs protein at both the plasma membrane and the Golgi apparatus in cardiomyocytes^23,33^. To test whether pharmacological manipulation of β1AR signaling can be utilized to resolve their compartmentalized function, we tested already existing β1AR antagonists possessing a range of hydrophobicity. Metoprolol, a hydrophobic/cell permeable β1AR antagonist, can inhibit both the plasma membrane and Golgi pools of β1ARs in HeLa cells^9^. In contrast, sotalol, a relatively hydrophilic βAR antagonist^34^ only inhibits the plasma membrane pool of β1AR in HeLa cells^9^. Following up on this logic, we then tested sotalol to isolate the role of Golgi-β1AR signaling in cardiomyocytes. We used our previously developed conformational sensitive nanobody-based biosensor, Nb80-GFP, to assess βARs activity in living cells^6,9^. We infected primary neonatal cardiomyocytes (NCM) with lentiviral constructs expressing β1AR and Nb80-GFP. In unstimulated β1AR expressing NCMs, Nb80-GFP expresses diffusely in the cytoplasm (Figure 1a, top row). Upon treatment of NCMs with 10 μM dobutamine, a cell permeable β1AR agonist, Nb80-GFP was recruited to activated β1AR at both the plasma membrane and the Golgi (Figure 1a, middle and bottom row; and Supplementary Movie 1).

**Figure 1:**
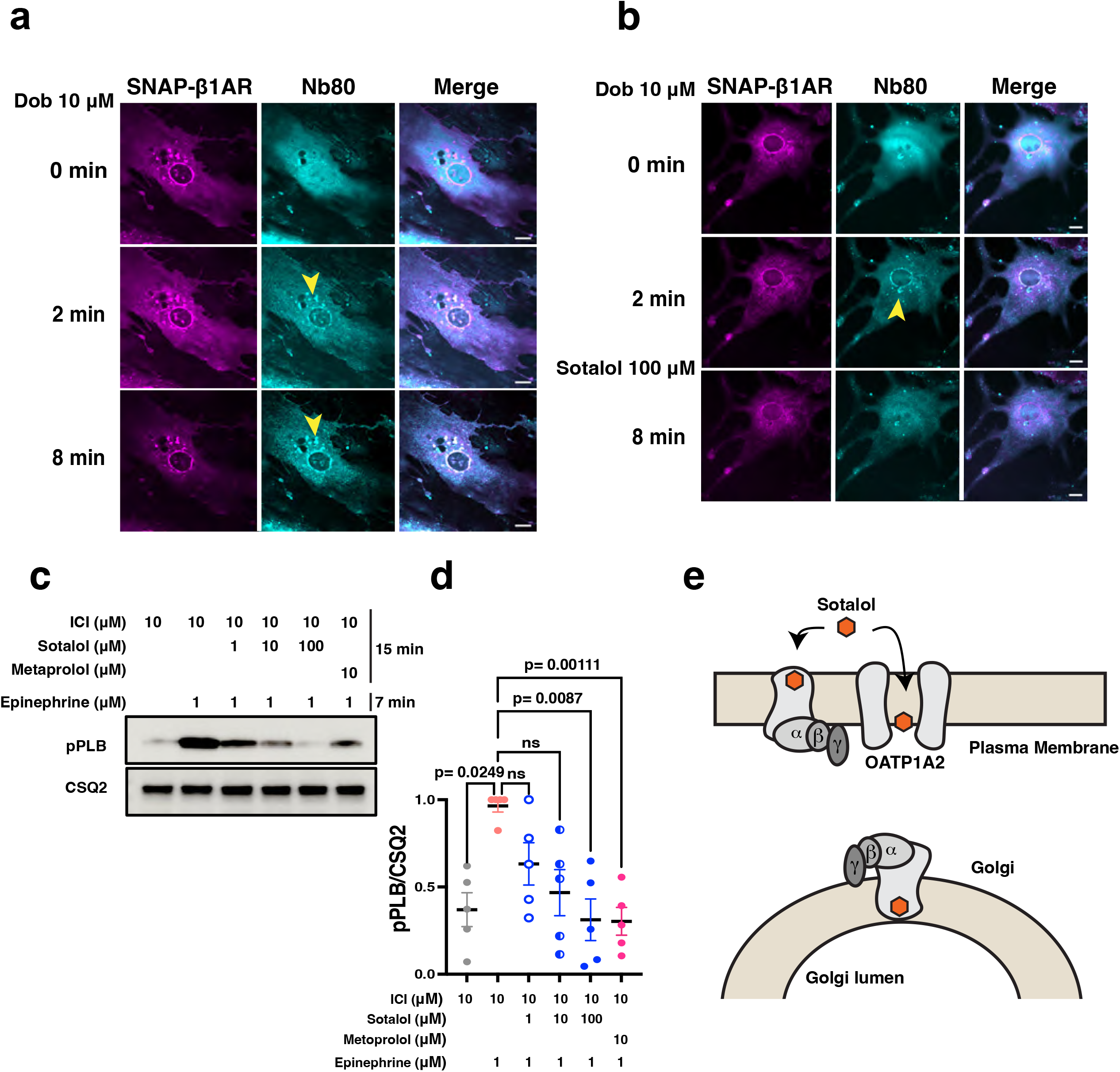
Sotalol inhibits both the plasma membrane and the Golgi-localized β1AR in cardiomyocytes. **a.** Representative confocal images of neonatal cardiomyocytes expressing the conformational biosensor of β1AR, Nb80-GFP (cyan), and SNAP-β1AR (magenta) before and after dobutamine (10 μM) treatment for 10 min. Stimulation with dobutamine (10 μM) results in Nb80-GFP recruitment to the plasma membrane and the Golgi. Arrowheads indicate Nb80-GFP localization at the Golgi. Scale bar: 10 μm. **b.** Representative confocal images of neonatal cardiomyocytes expressing Nb80-GFP (cyan) and SNAP-β1AR (magenta) pre-treated with dobutamine (10 μM) and before and after sotalol (100 μM) addition. Sotalol (100 μM) treatment for 3 min results in the loss of Nb80-GFP localization at the Golgi. Arrowheads indicate Nb80-GFP localization at the Golgi. Scale bar: 10 μm. **c.** Representative phosphorylation profile of PLB (pPLB) regulated by β1AR in adult cardiomyocytes derived from wild-type mice. Adult cardiomyocytes were pretreated with 10 μM β2AR-selective antagonist ICI-118551 (ICI) to isolate the function of β1AR. Phosphorylation of PLB Ser16/Thr17 (pPLB) were analyzed in wild-type adult cardiomyocytes (ACM) upon treatment with metoprolol (10 μM) or sotalol (1, 10 and 100 μM) for 15 min and followed by epinephrine (1 μM) treatment for 7 min at 37℃. **d.** Quantification of immunoblots of pPLB were normalized to the protein levels of CSQ2, and then reported as a percentage of the highest value in the groups. The quantified data from different experiments are presented as mean ± S.E.M. The *p*-values were calculated by two-way ANOVA. n = 5 independent biological replicates. **e.** Illustration of the facilitated transport of sotalol by OATP1A2. Sotalol (orange) can be transported by OATP1A2 intracellularly to inhibit Golgi-localized β1AR.

Having validated the response of Nb80-GFP in NCMs, we then tested the capability of sotalol in blocking the plasma membrane pool of β1AR. NCM expressing β1AR and Nb80-GFP were treated with 10 μM dobutamine and as expected we observed recruitment of Nb80-GFP in both PM and Golgi compartments (Figure 1b, upper and middle panel and Supplementary movie 2). To our surprise, however, addition of 100 μM sotalol resulted in inhibition of both pools of β1AR, as indicated by loss of Nb80-GFP localization at the plasma membrane and the Golgi (Figure 1b, lower panel). These results suggested that sotalol can reach the Golgi membrane and inhibit Golgi-localized β1AR.

To further confirm this result, we next assessed the effect of sotalol in inhibiting downstream effectors of β1AR signaling. Epinephrine stimulation of β1AR promotes phosphorylation of Ryanodine 2 receptors (RyR2), cardiac Troponin I (TnI) and Phospholamban (PLB) through PKA activation^35^. We have previously established that PLB phosphorylation is mediated by activation of the Golgi-β1AR signaling^33^. Thus, to test the effect of sotalol in blocking Golgi-β1AR signaling, we assessed PLB phosphorylation in isolated adult cardiomyocytes (ACM). We first compared the efficacy of metoprolol and sotalol (Figure S1a). Given that metoprolol is a more potent β1AR antagonist (Figure S1a and b), we pre-treated ACMs with various concentration of sotalol (1,10 and 100 μM) and compared in to 10 μM metoprolol. We then stimulated ACMs with epinephrine, the endogenous full agonist of β1AR. The western blot analysis showed that 10 and 100 μM sotalol treatment significantly block PLB phosphorylation and to the same extend as 10 μM metoprolol (Figure 1c). Altogether, these data suggest that sotalol can enter cardiomyocytes to inhibit Golgi-localized β1AR-mediated PLB phosphorylation (Figure 1e).

One prediction would be that cardiomyocytes express transporters that can facilitate sotalol transport. It has been reported that solute carrier organic anion transporter 1A2 (OATP1A2) can transport anionic drugs such as sotalol across lipid bilayer^30,31^. We thus tested the expression of OATP1A2 and found that it is expressed in adult and neonatal cardiomyocytes (Figure S2a). To test whether sotalol can be transported by OATP1A2, we then treated ACMs with naringin, a selective inhibitor of OATP1A2^36,37^. Importantly, treatment of ACMs with 100 μM naringin blocked the effect of 10 μM sotalol on PLB phosphorylation (Figure S2b and c). Altogether, these data suggest that OATP1A2 is expressed in ACMs and can facilitate the transport sotalol into the cells to block Golgi-localized β1AR signaling.

### Design and synthesis of cell-permeable and impermeable spatially restricted β1AR Antagonists

To overcome the limitation of facilitated transport of anionic drugs such as sotalol by OATP1A2, we sought to design impermeable antagonists of β1AR, that cannot be transported into cells and can thus be used in multiple cell lines and model systems. To find a suitable β1AR antagonist that selectively inhibits the plasma membrane pool of the receptor, we analyzed the structure-activity relationships of existing antagonists. We found that multiple existing β-blockers (i.e. atenolol, metoprolol, and bisoprolol) share the same core pharmacophore but differ in their substituents in para-position of the aromatic ring (Figure 2a)^38,39^. Thus, we decided to attach a linker on the aromatic ring to conjugate cyanine5 (Cy5) or impermeable sulfo-cyanine5 (sulfo-Cy5), resulting in a pair of permeable and impermeable antagonist for β1AR (Figure 2b and c). The choice of a Cy5 dye with or without a sulfo-moiety provides not only a means to restrict membrane permeability but a fluorescent dye to follow both pharmacological agents. To do this, an alkyne intermediate was generated in three synthetic steps from tyramine and 4-pentynoic acid and was subsequently coupled with azido-functionalized fluorophores using click chemistry (Supplementary Information). To test, if these functionalized ligands are indeed binding β1AR as expected, we conducted docking studies with PDB 7BTS ^40^. We obtained a docking score of -10.581 for poses in which the pharmacophore is engaging the orthosteric binding site through the interaction with two key residues (D1138, F1218) and the conjugated fluorophore extends out of the binding pocket (Figure 2d and e, Figure S3). These data suggest that the conjugated cell permeable and impermeable antagonists can engage β1AR similar to previously reported β1AR antagonists.

**Figure 2:**
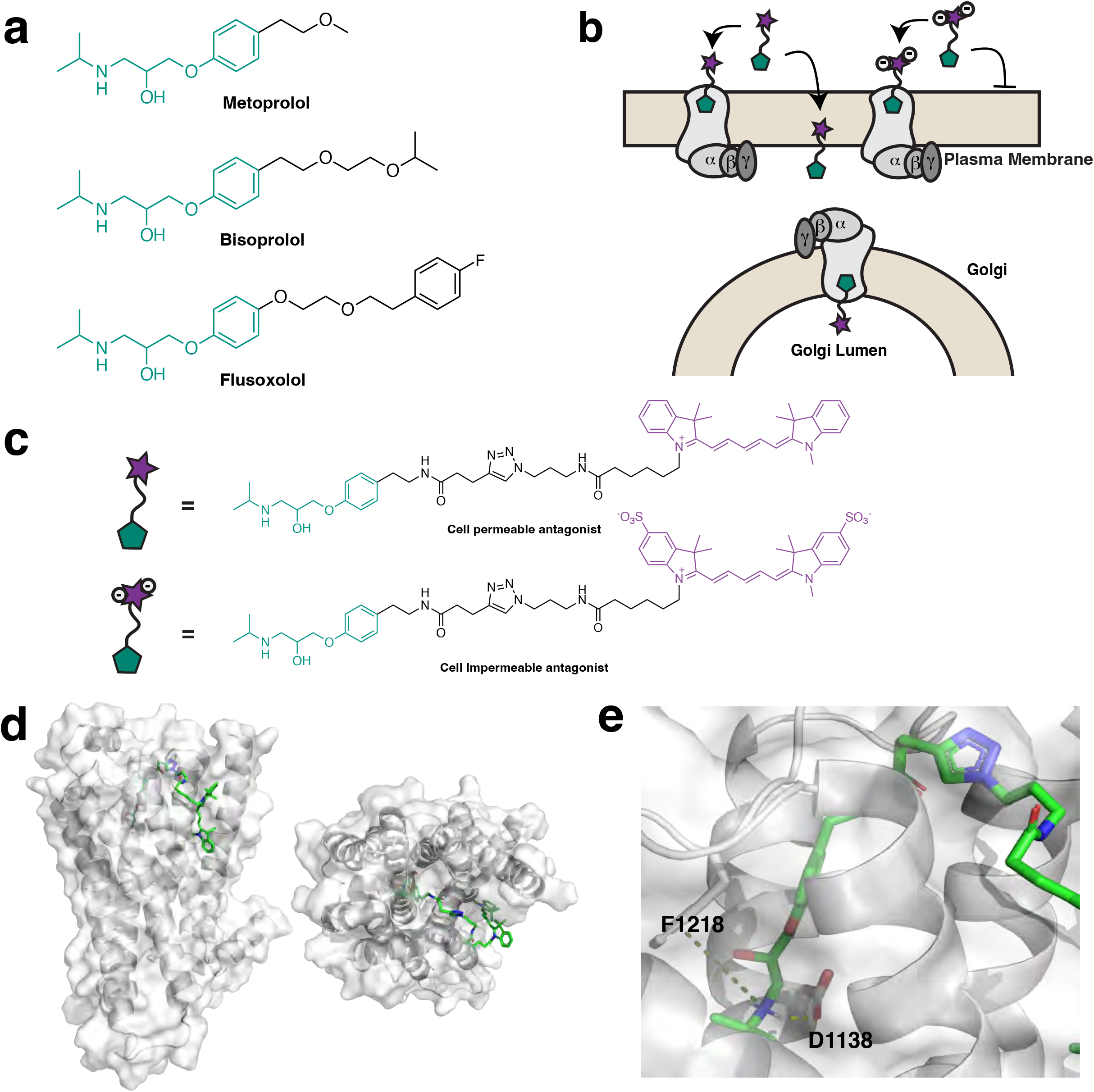
Rational design and synthesis of spatially restricted β1AR antagonists. **a.** Selection of β1AR antagonists as templates for the generation of spatially restricted antagonists. Common structural features of β1AR selective antagonist metoprolol, bisoprolol and flusoxolol are highlighted in green and have been used as templates for the design of a spatially restricted antagonist. **b.** Illustration of the mechanism of action of the cell permeable and impermeable antagonists. **c.** Chemical structures of cell permeable and impermeable antagonist. Pharmacophore moiety of the antagonist (green) is functionalized with a linker (black) and a fluorophore (purple). The cell permeable antagonist is conjugated with a Cy5 dye while the cell impermeable antagonist is conjugated with a sulfonated Cy5. **d.** Computational analysis of the binding interaction between cell permeable antagonist and β1AR (PDB:7BTS). **e.** β1AR orthosteric pocket and pharmacophore interaction sites. D1138 and F1218 are β1AR residues interacting with the pharmacophore via H-bond an ρε-cation interaction, respectively.

### Characterization of spatially restricted β1AR antagonists

To assess whether our newly designed β1AR antagonists’ function in a spatially restricted manner, we first checked their cellular distributions. Given that these antagonists are conjugated to a sulfonated or non-sulfonated Cy5, they are intrinsically fluorescent and thus can be visualized by live-imaging. We found that while cell permeable β1AR antagonist rapidly accumulates inside the cells, the cell impermeable antagonist is unable to cross the plasma membrane and remains outside the cells (Figure S4a and b). We next determined the pharmacological properties of these antagonists. Both agonists have similar efficacy and potency, as measured by their inhibitory effect on β1AR-mediated cAMP response (Figure 3a). Having determined the most effective concentrations (10 μM), we then tested their ability to selectively inhibit β1AR signaling, using our nanobody-based biosensor. HeLa cells expressing SNAP-tagged β1AR and Nb80-GFP were stimulated with 1 μM dobutamine for 15 min. Dobutamine is a cell permeable β1AR agonist and activates both the plasma membrane and the Golgi pool of β1AR, as demonstrated by Nb80-GFP recruitment to both locations (Figure 3b, upper panel). We then add 10 μM of the cell impermeable antagonist and noticed that while Nb80-GFP was still associated with the Golgi membranes, its localization to the plasma membrane was lost, suggesting that only the plasma membrane-β1AR is inhibited (Figure 3b, lower panel and c). In contrast, the cell permeable antagonist inhibits β1ARs at the plasma membrane and the Golgi and thus promotes Nb80-GFP dissociation from both compartments (Figure 3d and e). Given that both drugs have comparable potencies and efficacies, these data suggest that the sulfonated chemical group makes the antagonist cell impermeable, thus only inhibits plasma membrane-localized β1AR.

**Figure 3:**
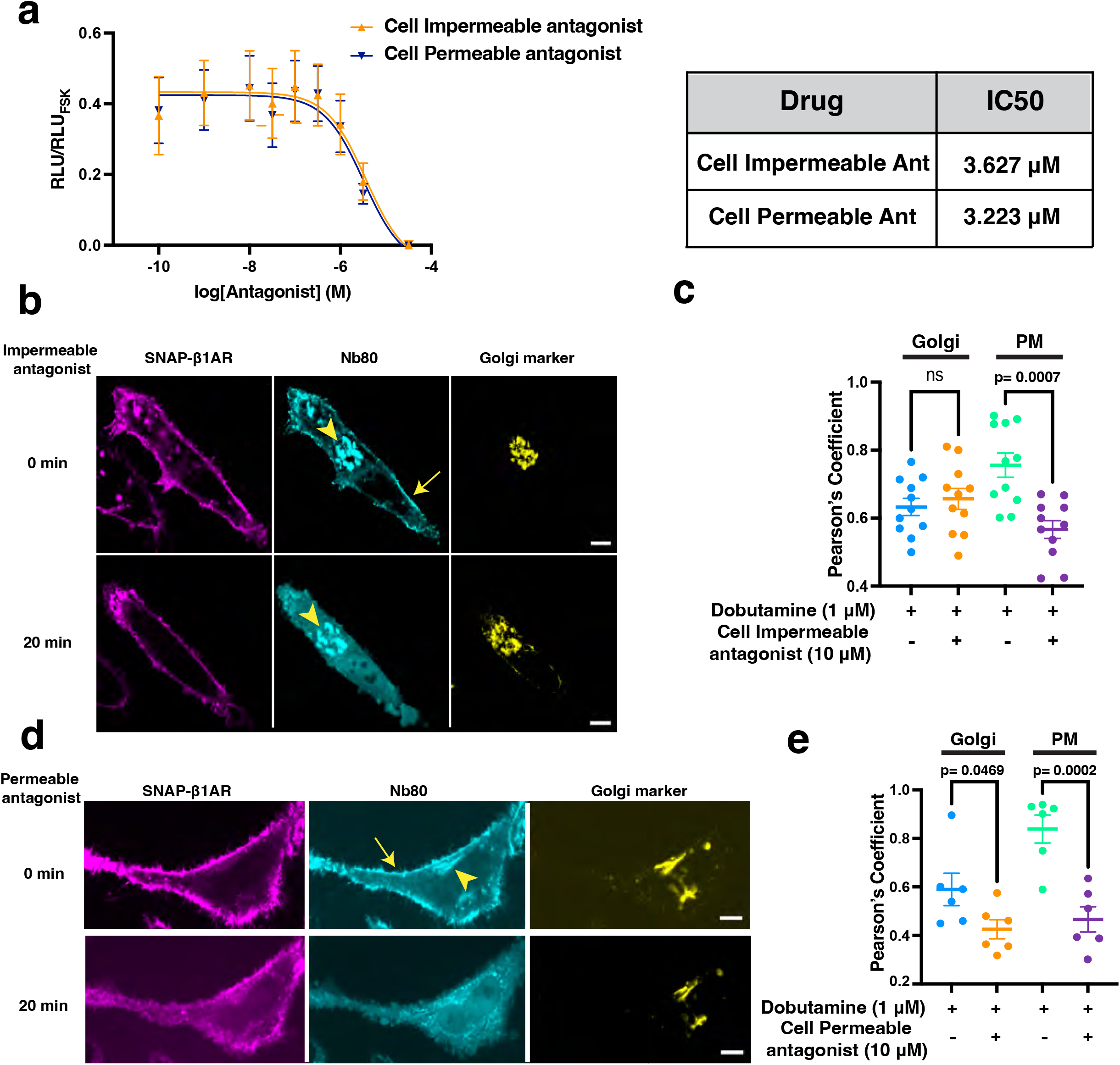
Conformational biosensor, Nb80-GFP, detects activated β1AR before and after cell permeable or impermeable antagonists’ treatment. **a.** Concentration-response curve of the cell permeable (blue line) and the cell impermeable (orange line) antagonists in HEK293 overexpressing β1AR and a bioluminescent cAMP biosensor. Cells were treated with different concentrations of antagonists in combination with ∼ 10 nM epinephrine. Relative luminescence values are normalized to HEK293 cells treated with forskolin (20 μM). n=4 biological replicates. IC50 values of cell impermeable and permeable antagonist are respectively 3.627 μM and 3.223 μM. **b.** Confocal images of representative HeLa cells expressing Nb80-GFP (cyan) and SNAP-β1AR (SNAP surface staining, magenta) and the Golgi marker (yellow), pre-treated with dobutamine (1 μM) for 15 min. Stimulation with cell impermeable antagonist (10 μM) for 20 min selectively inhibits the plasma membrane-localized β1AR. Arrow indicates plasma membrane and arrowheads indicate Golgi localization. Scale bar: 10 μm **c.** Person’s correlation coefficient measurements for the co-localization analysis of Nb80-GFP at the Golgi and plasma membrane after treatments of cardiomyocytes with the cell impermeable antagonist. The results have been measured from n=10 cells, 3 biological replicates; *p* values were calculated by one-way ANOVA. **d.** Confocal images of representative HeLa cells expressing Nb80-GFP (cyan) and SNAP-β1AR (SNAP surface staining, magenta) and the Golgi marker (yellow), pre-treated with dobutamine (1 μM) for 15 min. Stimulation with cell permeable antagonist (10 μM) for 20 min inhibits both the plasma membrane and the Golgi pool of β1AR. Arrow indicates plasma membrane and arrowheads indicate Golgi localization. Scale bar: 10 μm**. e.** Person’s correlation coefficient measurements for the co-localization analysis of Nb80-GFP at the Golgi and plasma membrane for cardiomyocytes treated with the cell permeable antagonist. The results have been measured from n=6 cells, 2 biological replicates; *p*-values were calculated by one-way ANOVA.

### Cell impermeable β1AR antagonist selectively block β1AR signaling on the plasma membrane in adult cardiomyocytes

In cardiomyocytes, β1AR-mediated cAMP generation regulates heart rate, force of contraction and relaxation through PKA-mediated phosphorylation of proteins, such as TnI, RyR2, and PLB ^35^. We have recently demonstrated that activation of the plasma membrane-localized β1ARs promote PKA-mediated phosphorylation of TnI and RyR2, respectively. In contrast, the Golgi-β1AR signaling specifically phosphorylates PLB^33^. To functionally test the compartment-specific inhibitory effects of our newly designed antagonists, we next assessed the phosphorylation status of downstream effectors of PKA upon β1AR activation. We specifically measured RyR2 and PLB phosphorylation, as readouts of the plasma membrane and Golgi-localized β1AR signaling, respectively. Treatment of ACMs with cell permeable β1AR antagonist resulted in the inhibition of both RyR2 and PLB phosphorylation upon epinephrine stimulations (Figure 4a and b). Importantly, treatment of ACMs with cell impermeable β1AR antagonist, only blocked RyR2 phosphorylation, while PLB phosphorylation was not affected (Figure 4 a and b). These data further confirm the selectivity effects of these compartment-specific antagonists.

**Figure 4:**
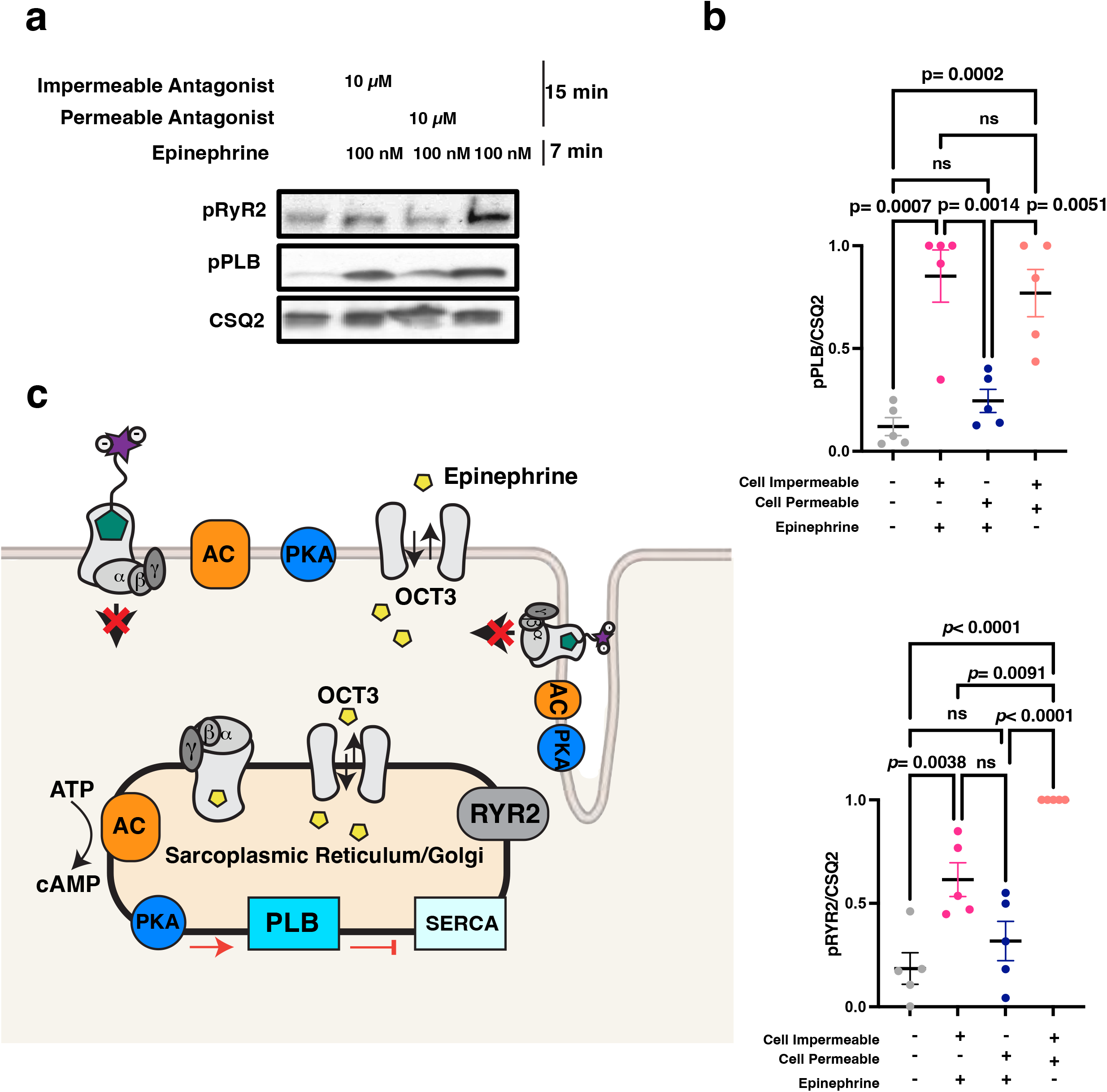
Cell impermeable antagonist selectively blocks plasma membrane-localized β1AR and inhibits RyR2 phosphorylation. **a.** Representative phosphorylation profiles of RyR2 and PLB induced by epinephrine (100nM) alone, or in combination with the cell impermeable and permeable antagonists (10 μM) in adult cardiomyocytes (ACM). Phosphorylation of RyR2 Ser2808 (pRyR2) and PLB Ser16/Thr17 (pPLB) were analyzed in wild-type ACMs endogenously expressing β1AR by immunoblotting. **b.** Quantification of immunoblots of pPLB and pRYR2 were normalized to the protein levels of CSQ2, and then reported as a percentage of the highest value in the groups. The quantified data from different experiments are presented as mean ± S.E.M. The *p*-values were calculated by two-way ANOVA. n = 5 biological replicates. **c.** Illustration of the effect of membrane impermeable antagonist in blocking β1AR-mediated signaling at the plasma membrane in adult cardiomyocytes while β1AR-mediated signaling at the Golgi is not affected.

## Discussion

In this study, we report the development of first-in-class β-blockers that are spatially restricted and can be used to manipulate β1AR compartmentalized signaling across cell types and tissues. By conjugating the pharmacophore of known β-blockers to a sulfonated fluorophore, we generated a membrane impermeable β1AR antagonist. By comparing the membrane impermeable β1AR antagonist to the cell permeable version of the same drug with similar potency and efficacy, we further show the functional significance of their effect in regulating β1AR-mediated signaling in cardiomyocytes. The cell-impermeable antagonist only inhibited β1AR and downstream effectors of PKA proximal to the plasma membrane while β1AR intracellular pool remained active demonstrated through continued phosphorylation of PLB. In contrast, cell permeable β1AR agonist efficiently inhibited both the plasma membrane and Golgi pools of β1ARs. We believe that this strategy allows better interrogation of compartmentalized β1AR signaling for *in vitro* and *in vivo* studies. It can also be potentially employed to manipulate several other GPCRs in a compartment specific manner.

In a recent study, a positive correlation between lipophilicity, based on the calculated logP values of chemically modified serotonin ligands, and their psychoplastogenic effects was reported^41^. The modification of GPCR agonists or antagonists with sulfonate groups has been previously used to visualize the distribution of receptors on the cell surface. For example, a sulfonated and fluorescently labelled analogue of an inverse βAR agonist, carazolol, or high-affinity agonist BI-167107 have been used to detect the distributions and dynamics of β1AR and β2AR on the plasma membrane^42,43^. However, none of these modified drugs were used to resolve the spatial functions of βARs. Our present study reports the first direct assessment of such chemical modifications and their consequent effects on cellular responses mediated by β1AR signaling.

Other approaches to manipulate intracellular GPCR signaling take advantage of chemically modified agonists and antagonists that are inactive until light-induced uncaging of active ligands ^44–48^. While this approach has been used to temporally control GPCR activity, spatial control has mainly been applied to map synaptic connectivity in neuronal networks^47^. There are few studies that have utilized caged compounds to manipulate spatial GPCR signaling at subcellular level. For example, a caged cell-permeable analog of the endothelin B antagonist was shown to specifically block intracellular endothelin receptors signaling^49^. Another report on angiotensin II receptors, took advantage of a caged version of Ang-II to demonstrate receptor signaling from intracellular compartments ^48^. We have previously used caged dopamine to activate D1DR signaling at the Golgi ^11^. While caged ligands can be useful to study subcellular GPCR activity in cells, they have some shortcomings for *in vivo* studies. For example, in some cases the side products of photolysis can create unintended consequences^47^. Moreover, many uncaged ligands can passively diffuse or be transported out of the cells via the same transport mechanism that facilitates their transport inside the cells. This is a particularly significant issue for caged monoamines (i.e., dopamine, epinephrine) that are transported by organic cation transporters (OCTs), known to function in a bidirectional manner^30^. Thus, uncaged monoamines can be transported in and out of the cells over time and activate both pools of monoamine GPCRs^11^. For example, we have previously demonstrated that epinephrine, an endogenous/hydrophilic β1AR agonist, requires a facilitated transport via an organic cation transporter, OCT3, to reach the β1ARs at the Golgi^9,23,33^. Similarly, we found that OCT2 facilitate the transport of dopamine to activate D1 dopamine receptor signaling at the Golgi^11^. Thus, in cells expressing OCT3 or OCT2, uncaged epinephrine or dopamine can be transported in and out of the cell via OCT3 or OCT2, respectively. This complexity makes the interpretation of the biological responses more complicated.

The complexity and variable tissue expression patterns of cation or anion transporters limits the utility of currently available GPCR drugs for studying their spatial function^30,32^. As reported in this study, sotalol, a relatively hydrophilic βAR antagonist can cross the membrane in a OATP1A2 dependent fashion. We found that HeLa cells do not express OATP1A2, thus sotalol cannot cross the membrane and only inhibit the plasma membrane-localized β1ARs. In contrast, adult cardiomyocytes express OATP1A2, thus sotalol can be transported into the cell and inhibit both the plasma membrane and the Golgi pool of β1ARs. The use of sulfonated GPCR agonist and antagonist particularly for monoamine GPCRs therefore offers the critical advantage to restrict drug access to subcellular compartment across cell types and tissues. The described strategy could be broadly applicable to modify other GPCR agonists and antagonists to resolve subcellular signaling profiles. It could further be useful in the clinical studies to distinguish between desired and undesired effects in therapy.

## Acknowledgments

We thank members of the Irannejad and the Shokat Labs for their assistance, advice, and valuable discussion. We also thank The Center for Advance Imaging (CALM) for helping with the biosensor imaging assay and analysis at UCSF. These studies were supported by the National Institute of General Medicine (GM133521 to R.I.), the Sandler Program for Breakthrough Biomedical research (PBBR7031249 to F.L), the National Cancer Institute (NCI) (K99CA277358 to J.M.), and (5R01CA244550 to K.M.S).

## Author contributions

F.L. designed the experimental strategy, carried out all western blot experiments and analysis, the viral preparation and the imaging experiments and helped the writing of the manuscript. J.M. performed all the chemical synthesis and analysis, the HPLC assays and analysis and helped writing of the manuscript. T.L. carried out the adult cardiomyocytes’ preparation and experiment. J.P. performed the docking experiments and interpretation. K.M.S. contributed to the experimental strategy and data interpretation of the chemical synthesis and docking. R.I. together with F.L. and J.M. and K.M.S. designed the experimental strategy and contributed to interpreting the results and writing the manuscript.

## Figure legends

**Figure S1:**
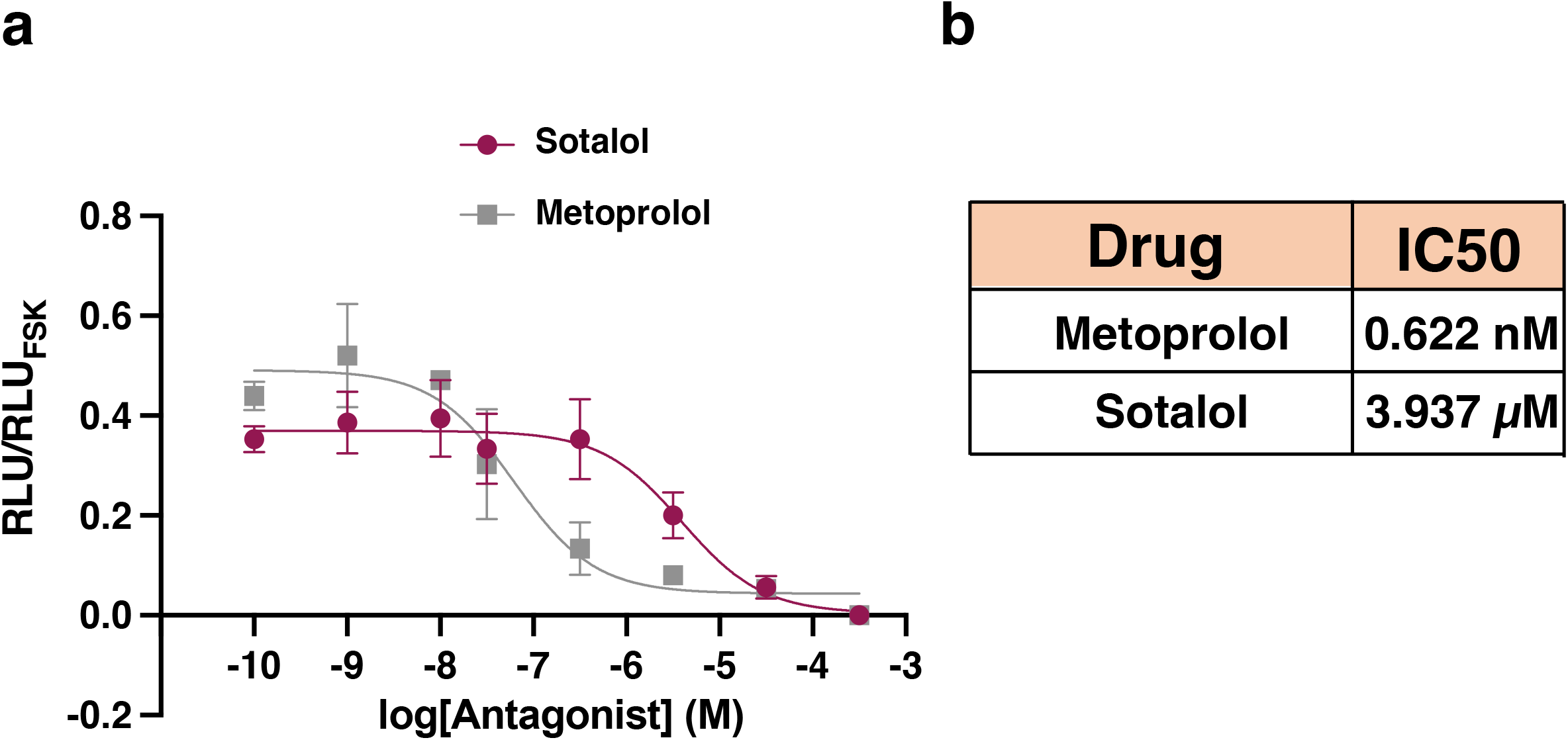
Concentration-response curve of sotalol and metoprolol. **a.** HEK293 overexpressing β1AR and the cAMP biosensor (pGlo Promega) were treated with various concentrations of antagonists. Concentration-response curve of sotalol (red) and metoprolol (grey) were obtained by normalizing the luminescence values of each condition with luminescence values obtained after treating cells with forskolin (20 μM). n=3 biological replicates. **b.** Calculated IC50 values of sotalol and metoprolol.

**Figure S2:**
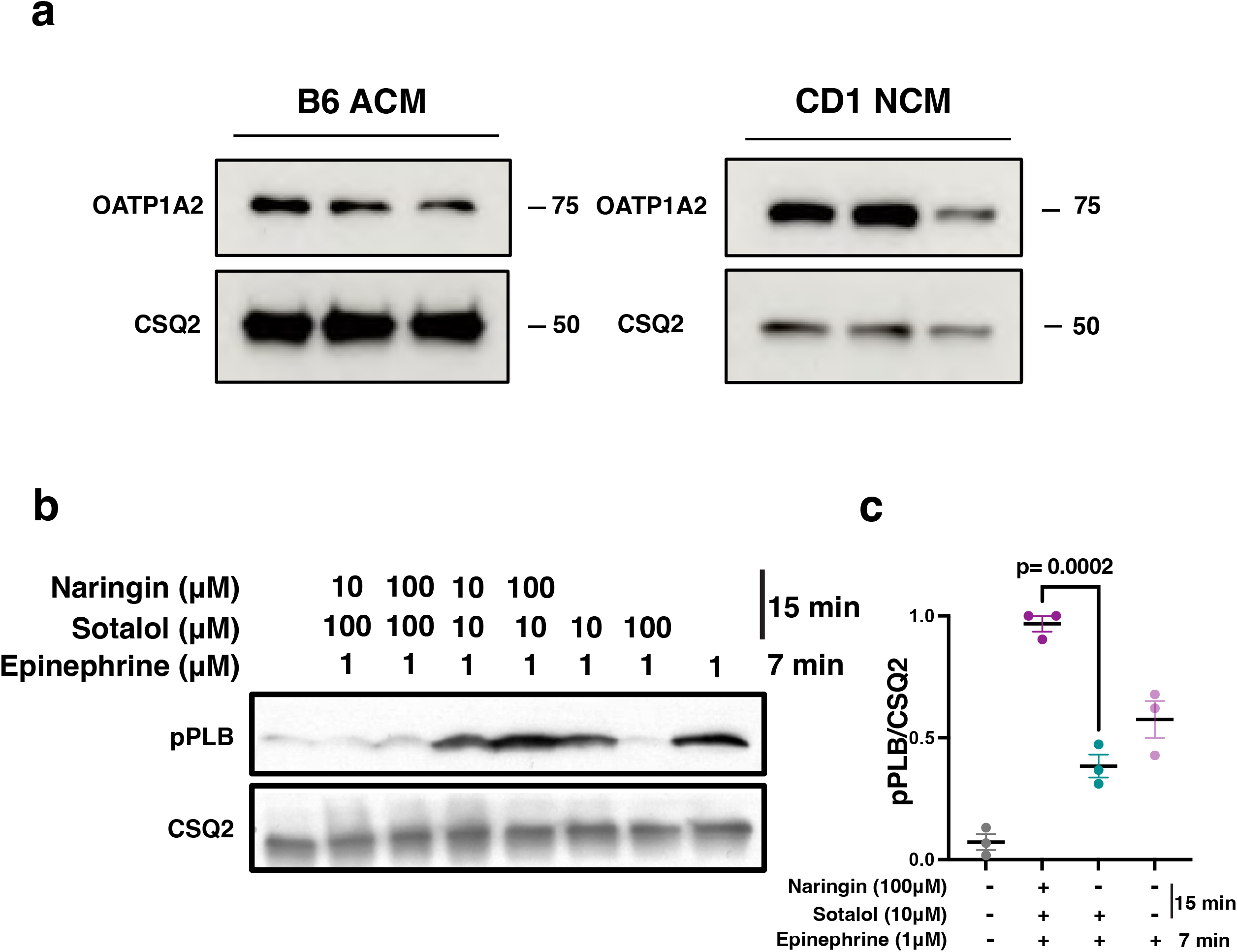
OATP1A2 is expressed in adult and neonatal murine cardiomyocytes. Representative western blots of OATP1A2 and CSQ2 expression in adult cardiomyocytes isolated from C57BL/6 (left) and neonatal cardiomyocytes from CD1 (right) mice. **b.** Representative phosphorylation profile of PLB (pPLB) regulated by β1AR in adult cardiomyocytes derived from wild-type mice. Adult cardiomyocytes were pretreated with 10 and 100 μM naringin for 15 min to inhibit OATP1A2. Phosphorylation of PLB Ser16/Thr17 (pPLB) were analyzed in wild-type adult cardiomyocytes (ACM) upon treatment with sotalol (10 and 100 μM) for 15 min and followed by epinephrine (1 μM) treatment for 7 min at 37℃. Naringin (100 μM) blocks the effect of sotalol (10 μM) inhibition on pPLB. **c.** Quantification of immunoblots of pPLB was normalized to the protein levels of CSQ2, and then reported as a percentage of the highest value in the groups. The quantified data from different experiments are presented as mean ± S.E.M. The *p*-values were calculated by two-way ANOVA. n = 3 biological replicates.

**Figure S3:**
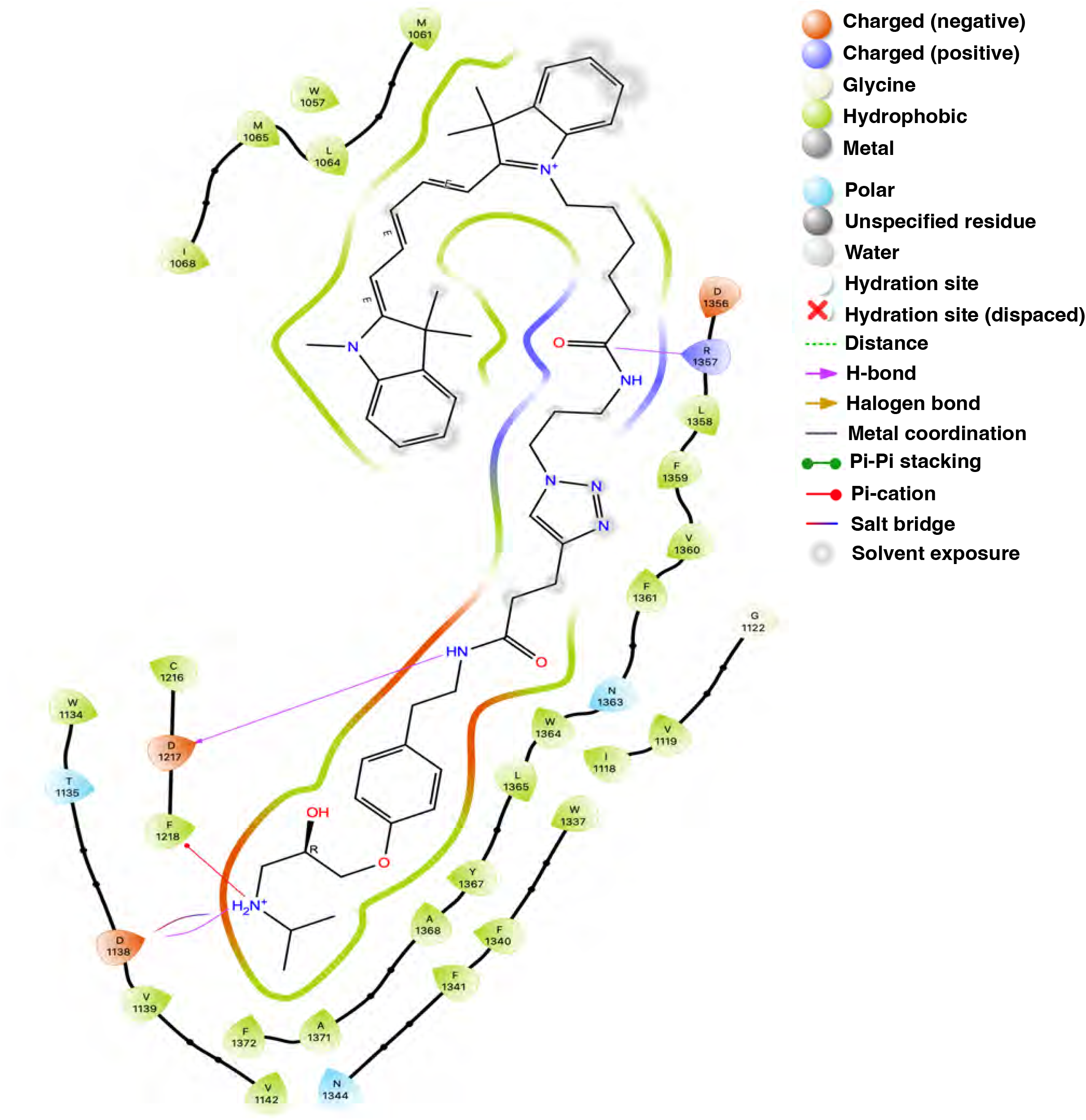
Computational analysis of β1AR cell permeable antagonist. Detailed analysis of cell permeable antagonist interaction sites with β1AR.

**Figure S4:**
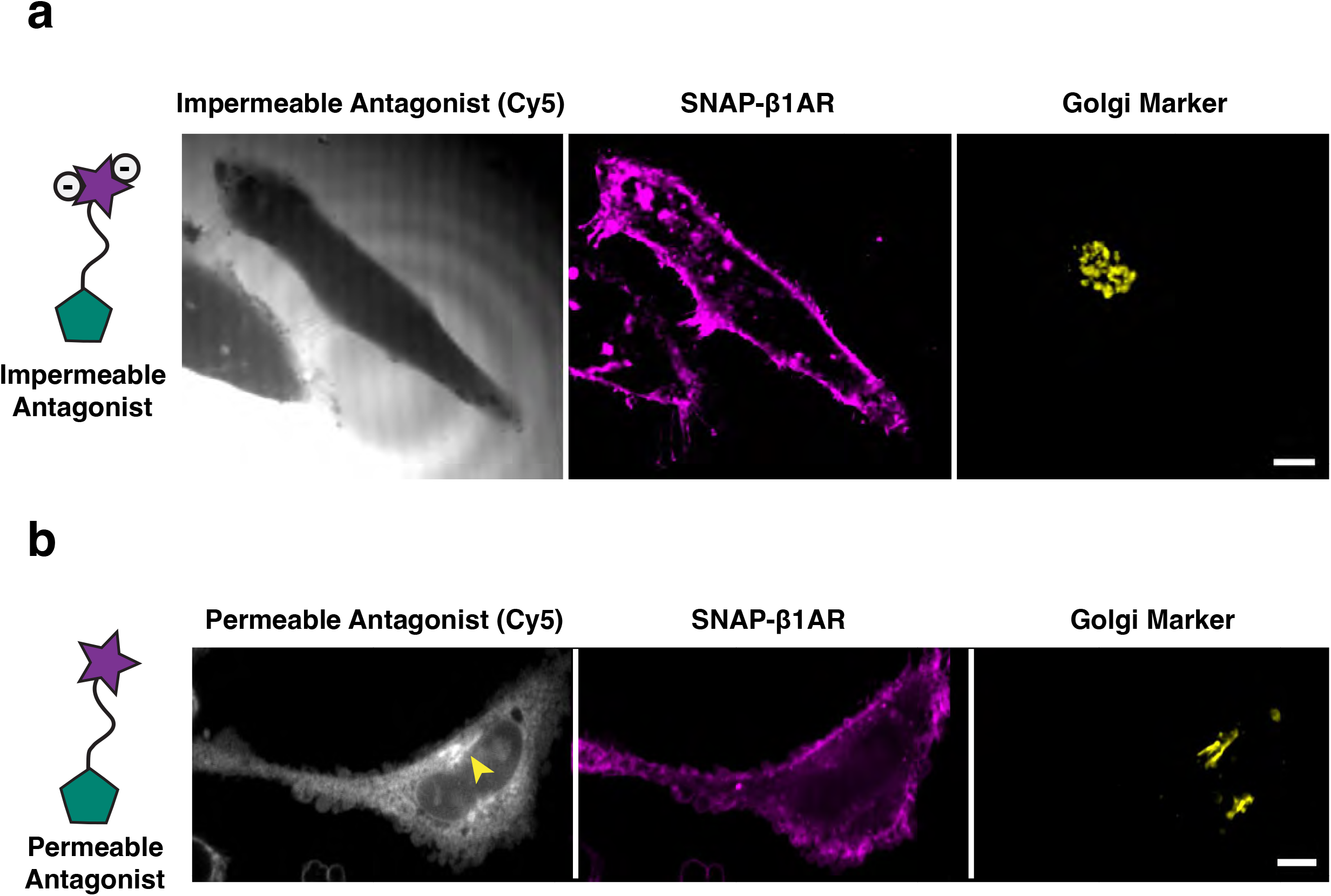
Cell impermeable antagonist cannot enter β1AR expressing HeLa cells. **a.** Representative confocal image of HeLa cell overexpressing SNAP-β1AR (magenta) and the Golgi marker (yellow), upon treatment with cell impermeable antagonist. Membrane impermeable antagonist can be monitored by fluorescent microscopy as the emission of the sulfonated Cy5 (grey) dye is in the far-red spectrum. Scale bar: 10 μm. **b.** Representative confocal image of HeLa cell overexpressing SNAP-β1AR (magenta) and the Golgi marker (yellow) upon treatment with cell permeable antagonist. Cell permeable antagonist can be monitored by fluorescent microscopy as the emission of the Cy5 (grey) dye is in the far-red spectrum. Cell permeable antagonist can reach the Golgi membrane after 20 min. Arrowhead indicates Golgi. Scale bar: 10 μm.

## Methods

### Reagents and antibodies

Human insulin, human transferrin and sodium selenite (ITS); urethane; 2,3-butanedione monoxime (BDM); Taurine; protease XIV; polybrene; forskolin (FSK); epinephrine; dobutamine; sotalol and metoprolol are from Sigma. Glutamax solution, penicillin and streptomycin, 4-(2-hydroxyethyl)-1-piperazineethanesulfonic acid (HEPES) buffer, DMEM, DMEM F12, M199 medium, Hanks’ Balanced Salt Solution (HBSS) buffer, ultrapure H2O, mouse laminin and Halt protease and phosphatase inhibitor cocktail are from Thermo Fisher Scientific. Fetal bovine serum (FBS) and Nu-Serum IV are from Corning. Glucose, sodium phosphate monobasic monohydrate, sodium chloride, potassium chloride, magnesium chloride hexahydrate, Tris–base, ethylenediaminetetraacetic acid (EDTA), dithiothreitol (DTT), dimethyl sulfoxide (DMSO), Tween-20 and HEPES are from Fisher Bioreagents. Doxycycline is from Takara. Calcium chloride is from Acros Organics. ICI-118551 is from Tocris Bioscience. Collagenase II is from Worthington. Bovine serum albumin (BSA) and dry milk powder are from Research Products International. SNAP-Surface Alexa Fluor 546 (S9132S), SNAP-Cell TMR-Star (S9105S), SNAP-Cell 647 SiR (S9102S) are from New England Biolabs. Heparin solution is from Fresenius Kabi. Triton X-100 is from Bio-Rad. Rabbit anti-phospho PLB (Ser16/Thr17) antibody (8496) and rabbit anti-phospho TnI (Ser23/Ser24) antibody (4004) are from Cell Signaling. Rabbit anti-phospho ryanodine receptor 2 (Ser2808) antibody (PA5-104444) is from Thermo Fisher Scientific. Rabbit anti-calsequestrin 2 (CSQ2) antibody (18422-1-AP) is from Proteintech and rabbit anti-OATP1A2 antibody (ab221804) is from Abcam. Amersham ECL donkey anti-rabbit IgG (NA934V), horseradish peroxidase (HRP)-linked whole antibodies were purchased from GE Healthcare Life Sciences.

### General method for spatially restricted antagonist synthesis

Anhydrous solvents were purchased from Acros Organics. Unless specified below, all chemical reagents were purchased from Sigma-Aldrich, Oakwood, Ambeed, or Chemscene. Analytical thin layer chromatography (TLC) was performed using aluminum plates pre-coated with silica gel (0.25-mm, 60-Å pore size, 230−400 mesh, Merck KGA) impregnated with a fluorescent indicator (254 nm). TLC plates were visualized by exposure to ultraviolet light (UV). Flash column chromatography was performed with Teledyne ISCO CombiFlash EZ Prep chromatography system, employing pre-packed silica gel cartridges (Teledyne ISCO RediSep). Proton nuclear magnetic resonance (1H NMR) spectra were recorded on Bruker AvanceIII HD instrument (400 MHz/100 MHz/376 MHz) at 23 °C operating with the Bruker Topspin 3.1. NMR spectra were processed using Mestrenova (version 14.1.2). Proton chemical shifts are expressed in parts per million (ppm, δ scale) and are referenced to residual protium in the NMR solvent (CHCl3: δ 7.26, MeOD: δ 3.31). Data are represented as follows: chemical shift, multiplicity (s = singlet, d = doublet, t = triplet, q = quartet, dd = doublet of doublets, dt = doublet of triplets, m = multiplet, br = broad, app = apparent), integration, and coupling constant (J) in Hertz (Hz). High-resolution mass spectra were obtained using a Waters Xevo G2-XS time-of-flight mass spectrometer operating with Waters MassLynx software (version 4.2). When LC-MS analysis of the reaction mixture is indicated in the procedure, it was performed as follows. An aliquot (1 µL) of the reaction mixture (or the organic phase of a mini-workup mixture) was diluted with 100 µL 1:1 acetonitrile/water. 1 µL of the diluted solution was injected onto a Waters Acquity UPLC BEH C18 1.7 µm column and eluted with a linear gradient of 5–95% acetonitrile/water (+0.1% formic acid) over 3.0 min. Chromatograms were recorded with a UV detector set at 254 nm and a time-of-flight mass spectrometer (Waters Xevo G2-XS).s

### Computational Modeling

The β1-adrenergic receptor structure from 7BTS was prepared using the Protein Preparation Wizard tool in Maestro (Schrödinger Suite 2023-1, Schrödinger, LLC, New York), and parts of the lysozyme were removed. The ligand was prepared using the LigPrep tool and docked into the protein using Glide. A truncated version without fluorophore was docked initially and then used as a reference ligand to dock the full ligand. The resulting docking scores were -7.658 for the truncated and -10.581 for the full ligand.

### Cell culture and lentivirus production

HeLa, HE293T and HEK 293 cells are cultured in DMEM (11965092) with 10% FBS. HEK293T were co-transfected with pMD2.G, pSPAX2 and Nb80-GFP and SNAP-β1AR plasmids using as transfection reagent TransIT-Lenti (mir6600, Mirus bio). Lentivirus was produced in DMEM containing 10% FBS and 1% BSA and then concentrated using the Lenti-X concentrator (Takera Bio).

### Animals

CD1 and C57BL/6 WT mice were housed in the UCSF facilities controlled by standardized environmental parameters such as 12h light/12h dark cycle in 7 d per week, humidity 30-70%, temperature 20-26°C and constant access to water and foods. All animal experiments were approved by the Institutional of Animal Care and Use Committee of the University of California (IACUC), San Francisco.

### Primary culture of neonatal cardiomyocytes

Hearts collected from P1–2 neonatal CD1 pups were cut into small pieces in ice-cold HBSS containing 20 mM HEPES. Heart pieces mixed with 225 IU ml−1 collagenase II were incubated on a rotator at 37 °C for 5 min. After several pipetting, the released cells in the buffer were collected by centrifuge at 500g for 5 min. The undigested heart tissues were digested again, as described above, until the undigested tissue became white, and the size do not decrease. The cells from each digestion were pooled together and resuspended in the neonatal cardiomyocyte culture media, which is DMEM (11995065) containing 10% FBS, 10% Nu-Serum IV, 10 mM HEPES, ITS, 10 mM Glutamax, penicillin and streptomycin. The isolated cells are passed through a 40 µm strainer and plated in a regular petri dish to remove the most of fibroblasts at 37 °C for 2 h. The fibroblast supernatant containing neonatal cardiomyocytes were collected and plated on mouse laminin-coated imaging dishes. For the virus transduction, lentivirus was mixed with the culture media with polybrene (8 µg ml−1). Virus was removed after 1-d transduction. The transduced neonatal cardiomyocytes were further treated with doxycycline for 3 d.

### Primary culture of adult cardiomyocytes

Adult cardiomyocytes were isolated from 2- to 3-month-old C57BL/6 WT male mice using the Langedorff-free method^50^. We intraperitoneally injected heparin solution into the mouse (5 U g−1). After 10 min, urethane, dissolved in 0.9% NaCl, was also intraperitoneally injected into the mouse (2 mg g−1). Once the mouse was anesthetized, we exposed the heart and the inferior vena cava was cut to release the blood. First, EDTA buffer (130 mM NaCl, 5 mM KCl, 0.5 mM NaH2PO4-H2O, 10 mM HEPES, 10 mM glucose, 10 mM BDM, 10 mM Taurine, in ultrapure H2O, pH 7.8) is injected into the right ventricle, and subsequently the aorta was clamped. The clamped heart was moved to the EDTA buffer-containing dish and then the EDTA buffer was injected into the left ventricle. Then, the clamped heart was moved to the perfusion buffer (130 mM NaCl, 5 mM KCl, 0.5 mM NaH2PO4–H2O, 10 mM HEPES, 10 mM glucose, 10 mM BDM, 10 mM Taurine, 1 mM MgCl2–6H2O, in ultrapure H2O, pH 7.8) containing dish and the perfusion buffer was injected into the left ventricle. The clamped heart was further moved to the digestion buffer (perfusion buffer with 1 mg ml^−1^ collagenase II and 0.05 mg ml^−1^ protease XIV) containing dish, and then the digestion buffer was injected in to left ventricle. After digestion, the heart was cut into small pieces and gently triturated to dissociate the cardiomyocytes. The digestion processes were stopped by adding stop buffer (perfusion buffer with 10% FBS), and the suspended cardiomyocytes were passed through a 100 µm strainer. The cardiomyocytes were enriched by gravity sedimentation and reintroduced calcium gradually. We resuspended the cardiomyocytes in plating media (M199 media with 5% FBS, 10 mM BDM, penicillin and streptomycin) and seeded on mouse laminin-coated wells at 37 °C for 1 h. After washing out the unattached cells by culture media (M199 media with 0.1% BSA, 10 mM BDM, penicillin and streptomycin and ITS), the cardiomyocytes were cultured in culture media for the drug treatment.

### Luminescence-based cAMP assay

HEK293 cells were transfected with β1AR and a cyclic-permuted luciferase reporter construct (pGloSensor-20F, Promega) and luminescence values were measured, as described previously^6^. Briefly, cells were plated in 96-well dishes (∼100,000 cells per well) in DMEM F12 without phenol red/no serum (21041-025, Thermofisher) and equilibrated to 37°C in the SpectraMax plate reader and luminescence was measured every 2 min for 20 min. Software was used to calculate integrated luminescence intensity and background subtraction. The agonist experiments have been carried out adding different concentration of epinephrine (1237000, Sigma) and dobutamine (D0676, Sigma). The antagonist experiments (cell permeable and impermeable spatially selective antagonists, sotalol (S0278, Sigma) and metoprolol (21041, Sigma) have been carried out incubating the cells at 37°C for 30 min. Cells were stimulated using the previously calculated EC80 of epinephrine (1237000). 20 μM forskolin (F6885, Sigma) was used as a reference value in each multi-well plate and for each experimental condition. The average luminescence value (measured across triplicate wells) was normalized to the maximum luminescence value measured in the presence of 20 μM forskolin (F6885, Sigma).

### Live-cell confocal imaging

Live-cell imaging was carried out using Nikon spinning disk confocal microscope with a ×60, 1.4 numerical aperture, oil objective and a CO2 and 37°C temperature-controlled incubator. A 406, 488, 568 and 640 nm Voltran was used as light sources for imaging mTagBFP2, GFP, SNAP-Surface Alexa Fluor 546, SNAP-Cell TMR-Star and SNAP-Cell 647 SiR signals, respectively. In the experiments conducted in HeLa cells, cells expressing SNAP-tagged receptor, Nb80-GFP and GalT-mTagBFP2 (Golgi marker) were imaged in 35 mm bottom glass imaging dishes. Receptors were surface labeled by addition of SNAP-Surface Alexa Fluor 546 (1:1000, New England Biolabs) to DMEM F12 without serum and phenol red (21041-025, Thermofisher) supplemented with 30 mM HEPES pH 7.4 (15630080, Thermofisher) for 10 min. Live-cell images where we tested the effect of the spatially restricted antagonist in HeLa cells were carried out by incubating the cells in 1 μM dobutamine (D0676, Sigma) at 37°C for 20 min before indicated spatially selective antagonists were added. Cells were imaged before and after 15 min from antagonist addition. Time-lapse images were acquired with a CMOS camera (Photometrics) driven by Nikon Imaging Software (NS Elements). In the experiments conducted in mouse neonatal cardiomyocytes (NCM), cells were isolated from neonatal CD1 mice and infected with pLVX-TetOne-SNAP-β1AR and pUBC-Nb80-GFP lentivirus as previously described. Cell expressing both SNAP-β1AR and Nb80-GFP were imaged in 35 mm bottom glass imaging dishes. Receptors were labelled both at the surface and intracellularly by addiction of the cell permeable SNAP-Cell 647 SiR ligand (1:1000, New England Biolabs) in DMEM F12 without serum and phenol red (21041-025, Thermofisher) supplemented with 30 mM HEPES pH 7.4 (15630080, Thermofisher) for 20 min. Live-cell images of NCM stimulated with dobutamine had been carried out by imaging cells before the addiction of 10 μM dobutamine. After 5 min from the addiction of dobutamine, 100 μM sotalol (S0278) was added to the cells. Cells were imaged every 10 seconds for 15 min. Time-lapse images were acquired with a CMOS camera (Photometrics) driven by Nikon Imaging Software (NS Elements).

### Image analysis and statistical analysis

Images were saved as 16-bit TIFF files. Quantitative image analysis was carried out on unprocessed images using ImageJ (http://rsb.info.nih.gov/ij). Analysis of Nb80-GFP colocalization analysis at the plasma membrane and Golgi in HeLa cells was estimated after background subtraction by calculating the Pearson’s coefficient between the indicated image channels with the plasma membrane marker (Surface SNAP-β1AR) and Golgi marker channel (GalT-mTagBFP2), using the colocalization the plugin for ImageJ JaCoP (17210054). For visual presentation (but not quantitative analysis), image series were processed using Kalman stack filter in ImageJ. P values are determined using two-way ANOVA calculated with Prism 10.0 software (GraphPad Software).

### Western blotting

After drug treatments, the cardiomyocytes adult mice were collected and lysed by radio-immunoprecipitation assay (RIPA) buffer containing inhibitors of proteases and phosphatases at 4 °C for 30 min on the tube rotator. Supernatants were collected after centrifuging at 4 °C for 10 min, and the protein amounts were determined by BCA assay (Sigma). The proteins were denatured by boiling for 10 min in the DTT-containing sample buffer and separated by 4–20% Mini-PROTEIN TGX gels (Bio-Rad) and then transferred to the 0.2 µm polyvinylidene difluoride (PVDF) membrane (Bio-Rad). PVDF membrane was further blocked by TBST (TBS buffer with 0.1% Tween-20) containing 3% milk at room temperature for 1 h and then incubated with the primary antibody in TBST containing 5% BSA at 4 °C for O/N. The PVDF membrane was washed by TBST three times and then incubated with a secondary antibody diluted in TBST containing 3% milk at room temperature for 1 h. The unbonded secondary antibodies were removed by three times washing using TBST. The protein signals were visualized by ECL substrate (34580, Thermofisher). To evaluate the relative band intensities, we first scanned our films (300 ppi resolution) using an office scanner and convert them to an 8-bit format. We then inverted these images and subtracted the background using ImageJ software. The bands were selected using the rectangular selection tool on ImageJ. The relative band intensities were plot and measured; each peak was separated by straight-line selection tool. The area of each peak was measured by the Wand tool. Semi-quantified phosphoprotein bands were then normalized to the total lysate bands (CSQ2), and data were presented as fold change of the maximum value that we measured on each western blot. P values are determined using two-way ANOVA calculated with Prism 10.0 software (GraphPad Software).

## Supplementary Information

### Chemical Synthesis

Anhydrous solvents were purchased from Acros Organics. Unless specified below, all chemical reagents were purchased from Sigma-Aldrich, Oakwood, Ambeed, or Chemscene. Analytical thin layer chromatography (TLC) was performed using aluminum plates pre-coated with silica gel (0.25-mm, 60-Å pore size, 230−400 mesh, Merck KGA) impregnated with a fluorescent indicator (254 nm). TLC plates were visualized by exposure to ultraviolet light (UV). Flash column chromatography was performed with Teledyne ISCO CombiFlash EZ Prep chromatography system, employing pre-packed silica gel cartridges (Teledyne ISCO RediSep). Proton nuclear magnetic resonance (^1^H NMR) spectra were recorded on Bruker AvanceIII HD instrument (400 MHz) at 23 °C operating with the Bruker Topspin 3.1. NMR spectra were processed using Mestrenova (version 14.1.2). Proton chemical shifts are expressed in parts per million (ppm, δ scale) and are referenced to residual protium in the NMR solvent (CHCl_3_: δ 7.26). Data are represented as follows: chemical shift, multiplicity (s = singlet, d = doublet, t = triplet, q = quartet, dd = doublet of doublets, dt = doublet of triplets, m = multiplet, br = broad, app = apparent), integration, and coupling constant (J) in Hertz (Hz). High-resolution mass spectra were obtained using a Waters Xevo G2-XS time-of-flight mass spectrometer operating with Waters MassLynx software (version 4.2). When LC-MS analysis of the reaction mixture is indicated in the procedure, it was performed as follows. An aliquot (1 µL) of the reaction mixture (or the organic phase of a mini-workup mixture) was diluted with 100 µL 1:1 acetonitrile/water. 1 µL of the diluted solution was injected onto a Waters Acquity UPLC BEH C18 1.7 µm column and eluted with a linear gradient of 5–95% acetonitrile/water (+0.1% formic acid) over 3.0 min. Chromatograms were recorded with a UV detector set at 254 nm and a time-of-flight mass spectrometer (Waters Xevo G2-XS).

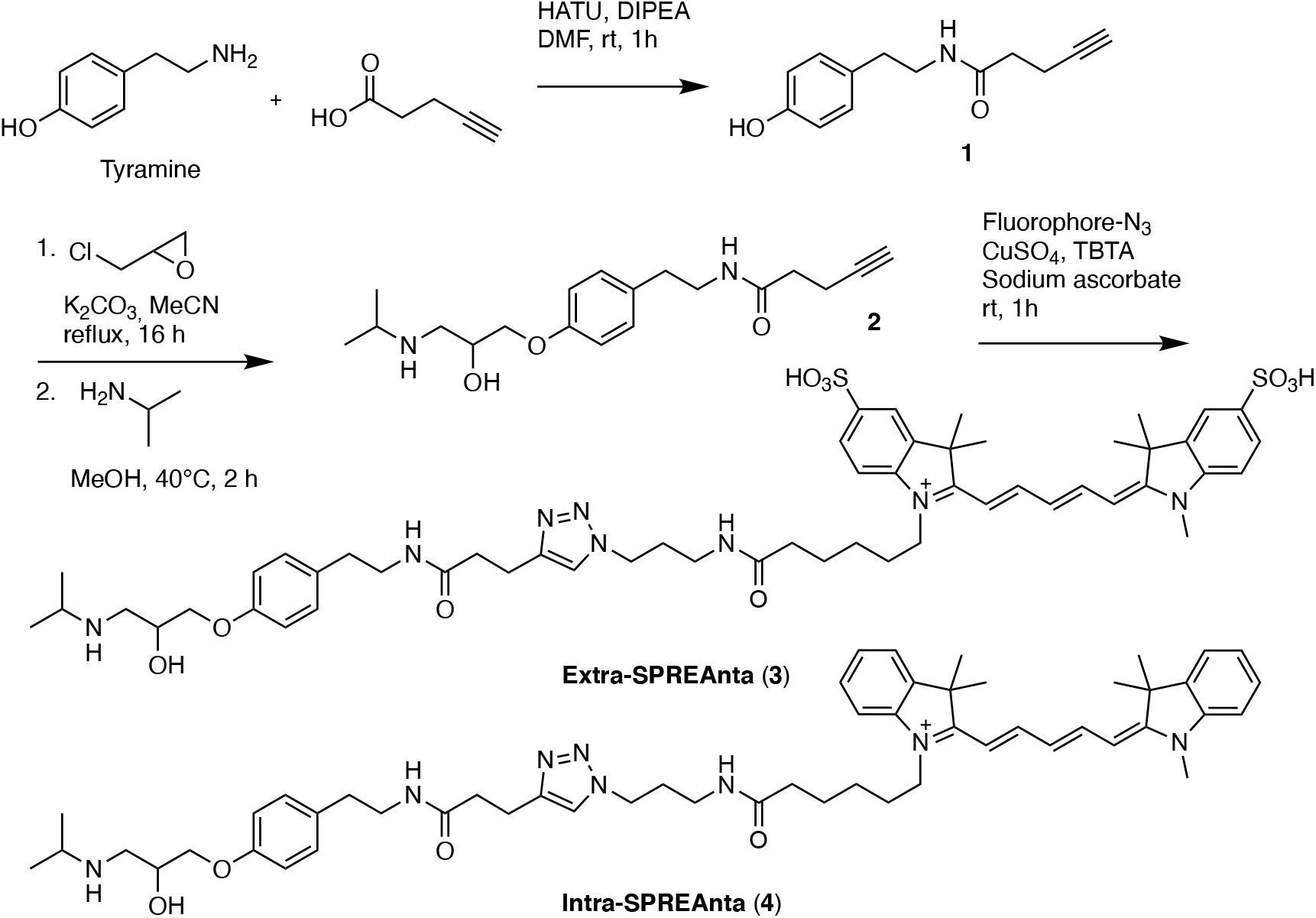

Scheme for the synthesis of **cell impermeable antagonist** (**3**) and cell **permeable antagonist** (**4**).

### Synthetic Details

#### *N-(4-hydroxyphenethyl)pent-4-ynamide* (1)

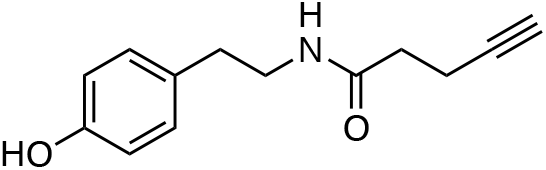

4-pentynoic acid (186 mg, 1.90 mmol, 1.3 equiv.), HATU (721 mg, 1.90 mmol, 1.3 equiv.), and DIPEA (565 mg, 4.37 mmol, 3.0 equiv.) were dissolved in DMF (5 mL) and stirred at room temperature for 10 min. Tyramine (200 mg, 1.46 mmol, 1.0 equiv.) was added and stirred at room temperature for 1 h. EtOAc and water were added, extracted with EtOAc and the combined organic layer was dried over NaSO_4_, filtered, and concentrated. The mixture was purified by flash column chromatography (20% EtOAc in hexanes to 100% EtOAc) to yield **1** as colorless glue (189 mg, 0.870 mmol, 60%).

**^1^H NMR** (400 MHz, CDCl_3_) δ 7.03 (d, *J* = 8.4 Hz, 2H), 6.83 – 6.75 (m, 2H), 6.49 (s, 1H), 5-75 (s, 1H), 3.50 (q, *J* = 6.9 Hz, 2H), 2.74 (t, *J* = 6.9 Hz, 2H), 2.49 (td, *J* = 6.9, 2.6 Hz, 2H), 2.35 (t, *J* = 7.1 Hz, 2H), 1.95 (s, 1H).

**LCMS**: m/z calcd. for C_13_H_16_NO_2_^+^ ([M]^+^): 218.1 found: 218.1.

#### *N-(4-(2-hydroxy-3-(isopropylamino)propoxy)phenethyl)pent-4-ynamide* (2)

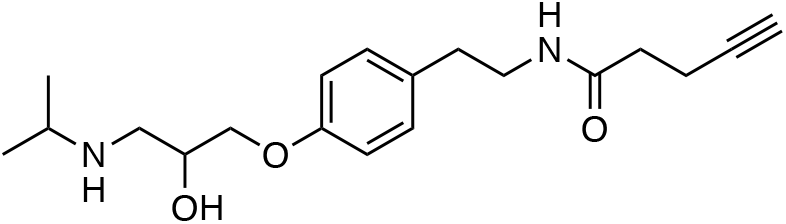

N-(4-hydroxyphenethyl)pent-4-ynamide (**1**) (189 mg, 0.870 mmol, 1.0 equiv.), (±)-epichlorohydrin (402 mg, 4.35 mmol, 5.0 equiv.), and K_2_CO_3_ (180 mg, 1.31 mmol, 1.5 equiv.) were added to MeCN (4 mL) and stirred under reflux for 16 h. The solvent was removed under reduced pressure and the crude was directly purified by flash column chromatography (40% EtOAc in hexanes to 100% EtOAc) to yield the intermediate as a white solid. The intermediate product was taken up in MeOH (3 ml) and 2-aminopropane (161 mg, 2.73 mmol, 5.0 equiv.) was added. The mixture was heated under reflux for 2 h and purified by reverse phase HPLC to yield **2** as a colorless solid (175 mg, 0.526 mmol, 60%).

**^1^H NMR** (400 MHz, CDCl_3_) δ 7.08 (d, *J* = 8.3 Hz, 2H), 6.80 (d, *J* = 8.4 Hz, 2H), 4.41 (s, 1H), 3.99 (d, *J* = 27.3 Hz, 2H), 3.48 (q, *J* = 6.7 Hz, 2H), 3.40 (s, 1H), 3.22 (s, 2H), 2.75 (t, *J* = 6.9 Hz, 2H), 2.49 (td, *J* = 6.9, 2.3 Hz, 2H), 2.36 (t, *J* = 7.1 Hz, 2H), 1.96 (d, *J* = 5.1 Hz, 1H), 1.39 (dd, *J* = 6.0, 3.8 Hz, 6H).

**LCMS**: m/z calcd. for C_19_H_29_N_2_O_3_^+^ ([M]^+^): 333.2 found: 333.2.

#### *1-(6-((3-(4-(3-((4-(2-hydroxy-3-(isopropylamino)propoxy)phenethyl)-amino)-3-oxopropyl)-1H-1,2,3-triazol-1-yl)propyl)amino)-6-oxohexyl)-3,3-dimethyl-2-((1E,3E)-5-((E)-1,3,3-trimethyl-5-sulfonatoindolin-2-ylidene)penta-1,3-dien-1-yl)-3H-indol-1-ium-5-sulfonate* (3)

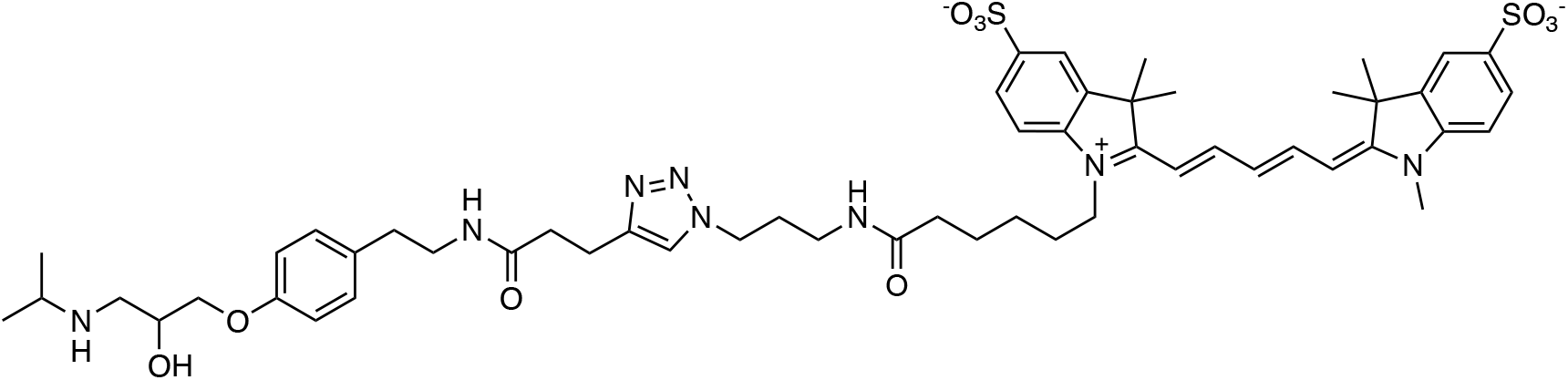

200 mL of a 20 mM solution of CuSO_4_ in water, 200 mL of a 500 mM solution of sodium ascorbate in water, and 200 mL of a 2 mM solution of TBTA in DMSO: butyl alcohol (1:4) were added to 200 mL of DMSO. Sulfo-Cy5 Azide (5 mg) was dissolved in 200 mL of DMSO, added, and stirred for 5 min at room temperature. **2** was dissolved in 200 mL of MeOH, added, and stirred at room temperature for 30 min. LCMS indicated full conversion to **3** and was purified by reverse-phase HPLC.

**HRMS**: m/z calcd. for C_54_H_73_N_8_O_10_^+^ ([M+H]^+^): 1057.4886 found: 1057.4874.

**HPLC**:

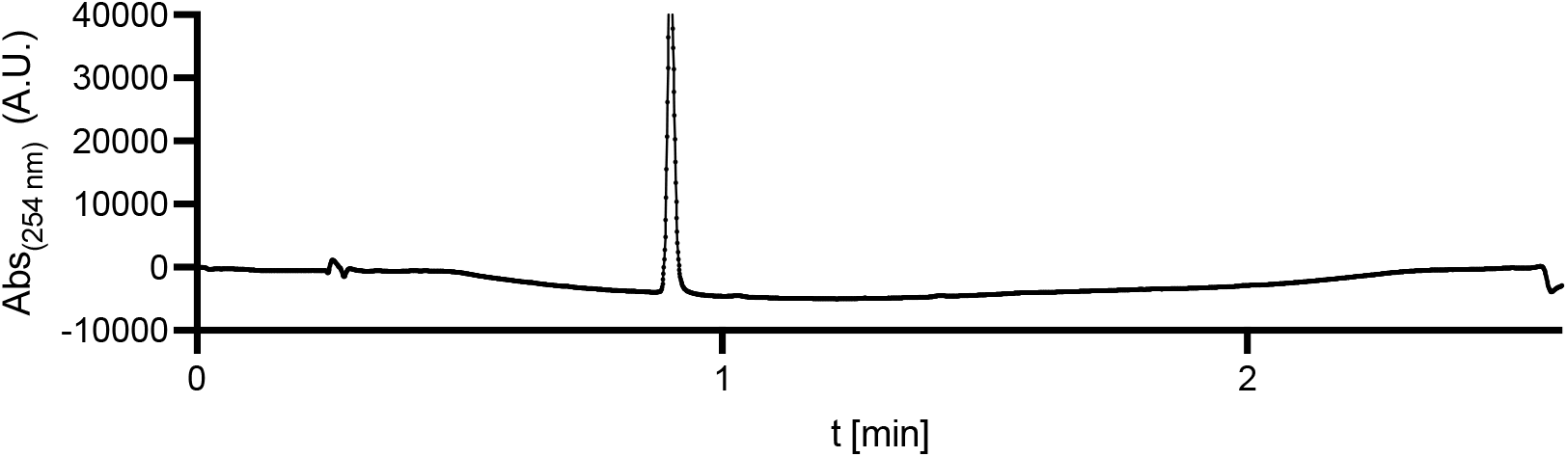

#### *1-(6-((3-(4-(3-((4-(2-hydroxy-3-(isopropylamino)propoxy)phen-ethyl)amino)-3-oxopropyl)-1H-1,2,3-triazol-1-yl)propyl)amino)-6-oxohexyl)-3,3-dimethyl-2-((1E,3E)-5-((E)-1,3,3-trimethylindolin-2-ylidene)penta-1,3-dien-1-yl)-3H-indol-1-ium* (4)

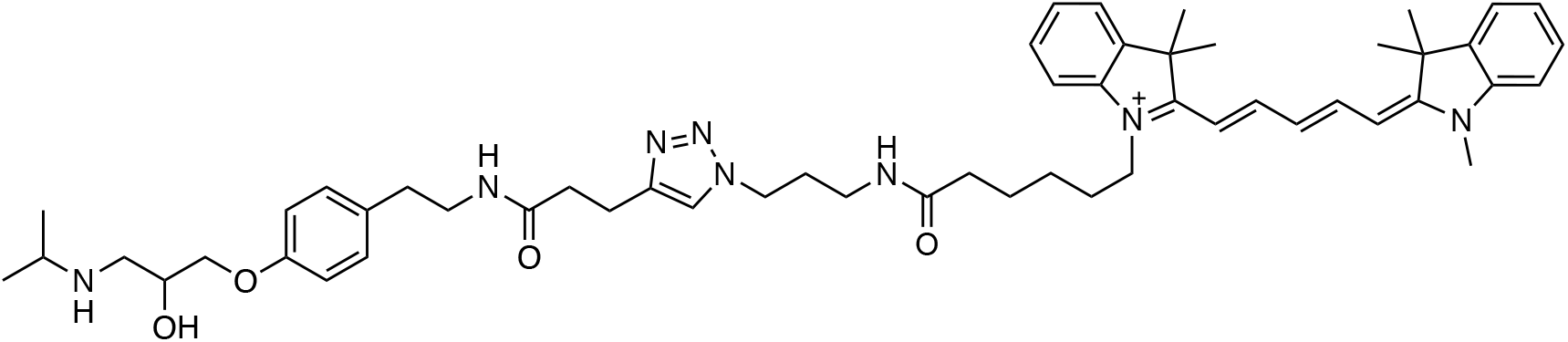

200 mL of a 20 mM solution of CuSO_4_ in water, 200 mL of a 500 mM solution of sodium ascorbate in water, and 200 mL of a 2 mM solution of TBTA in DMSO : butyl alcohol (1:4) were added to 200 mL of DMSO. Sulfo-Cy5 Azide (5 mg) was dissolved in 200 mL of DMSO, added, and stirred for 5 min at room temperature. **2** was dissolved in 200 mL of MeOH, added, and stirred at room temperature for 30 min. LCMS indicated full conversion to **4** and was purified by reverse-phase HPLC.

**HRMS**: m/z calcd. for C_54_H_73_N_8_O_4_^+^ ([M+H]^+^): 897.5749 found: 897.5756.

**HPLC:**

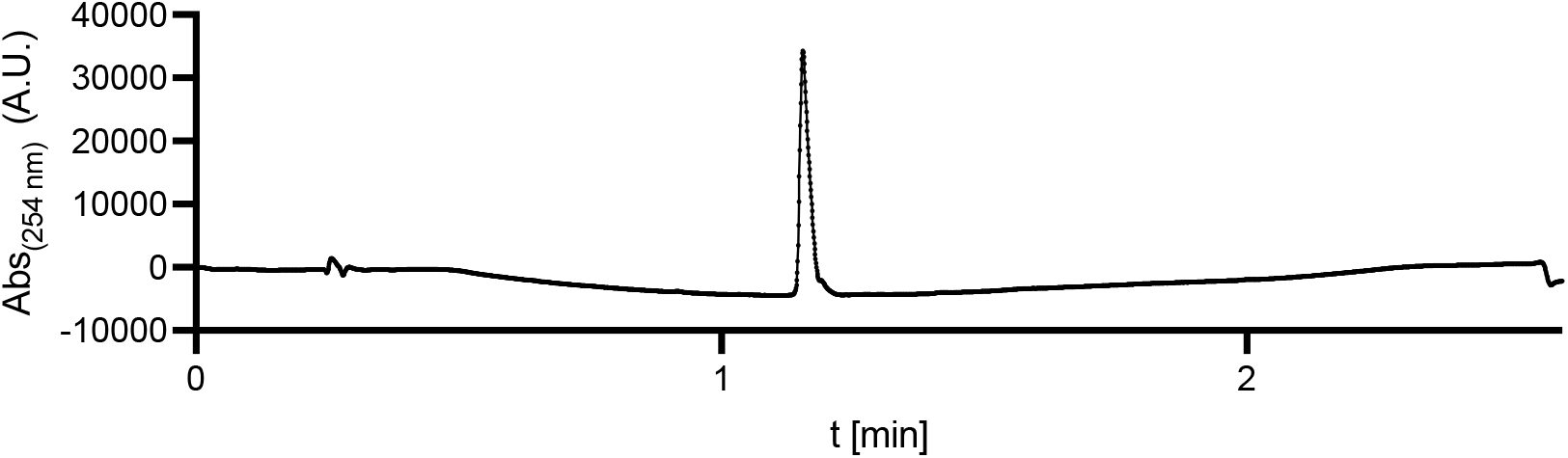

### NMR Spectra

#### *N-(4-hydroxyphenethyl)pent-4-ynamide* (1)

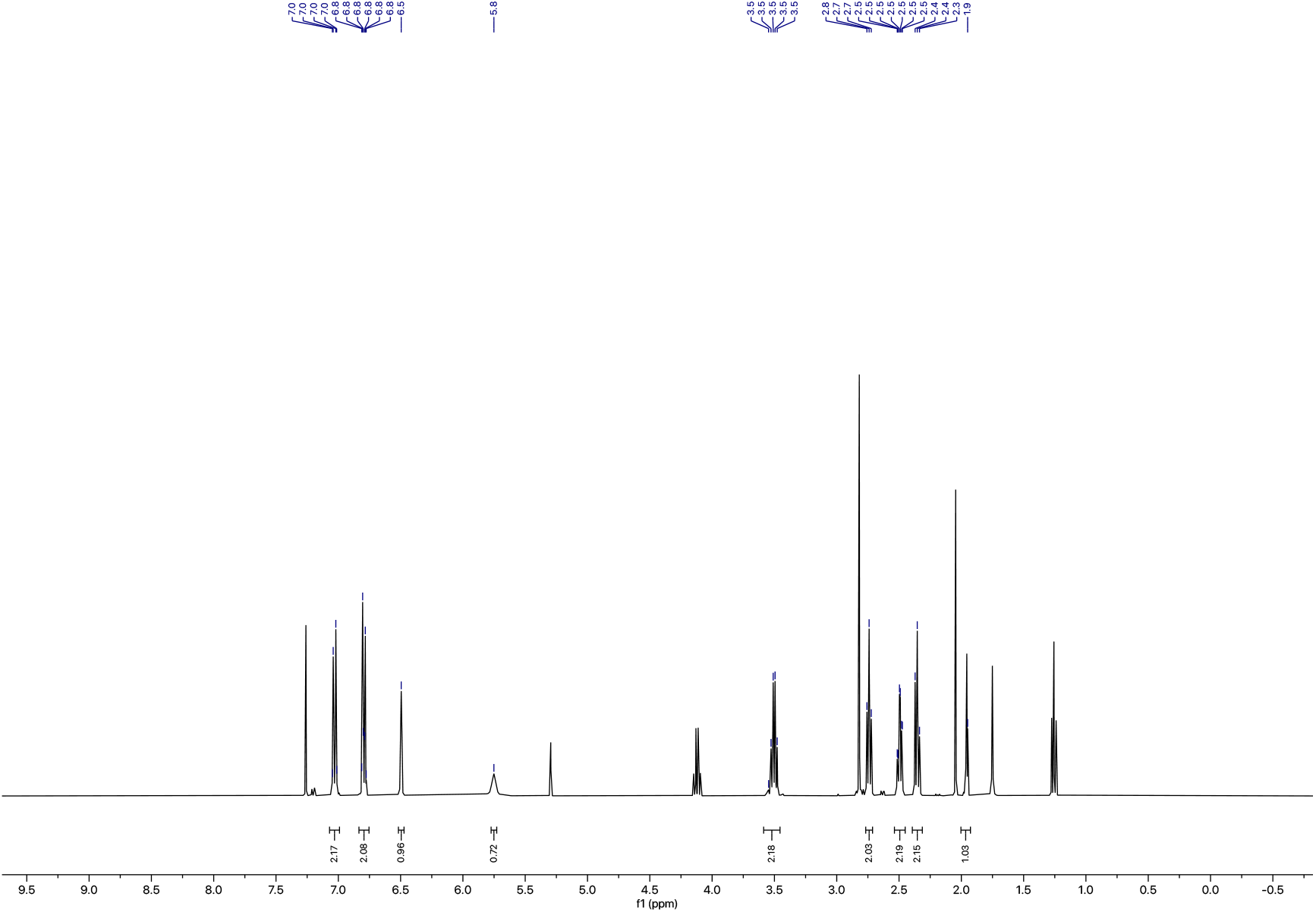

#### *N-(4-(2-hydroxy-3-(isopropylamino) propoxy)phenethyl)pent-4-ynamide* (2)

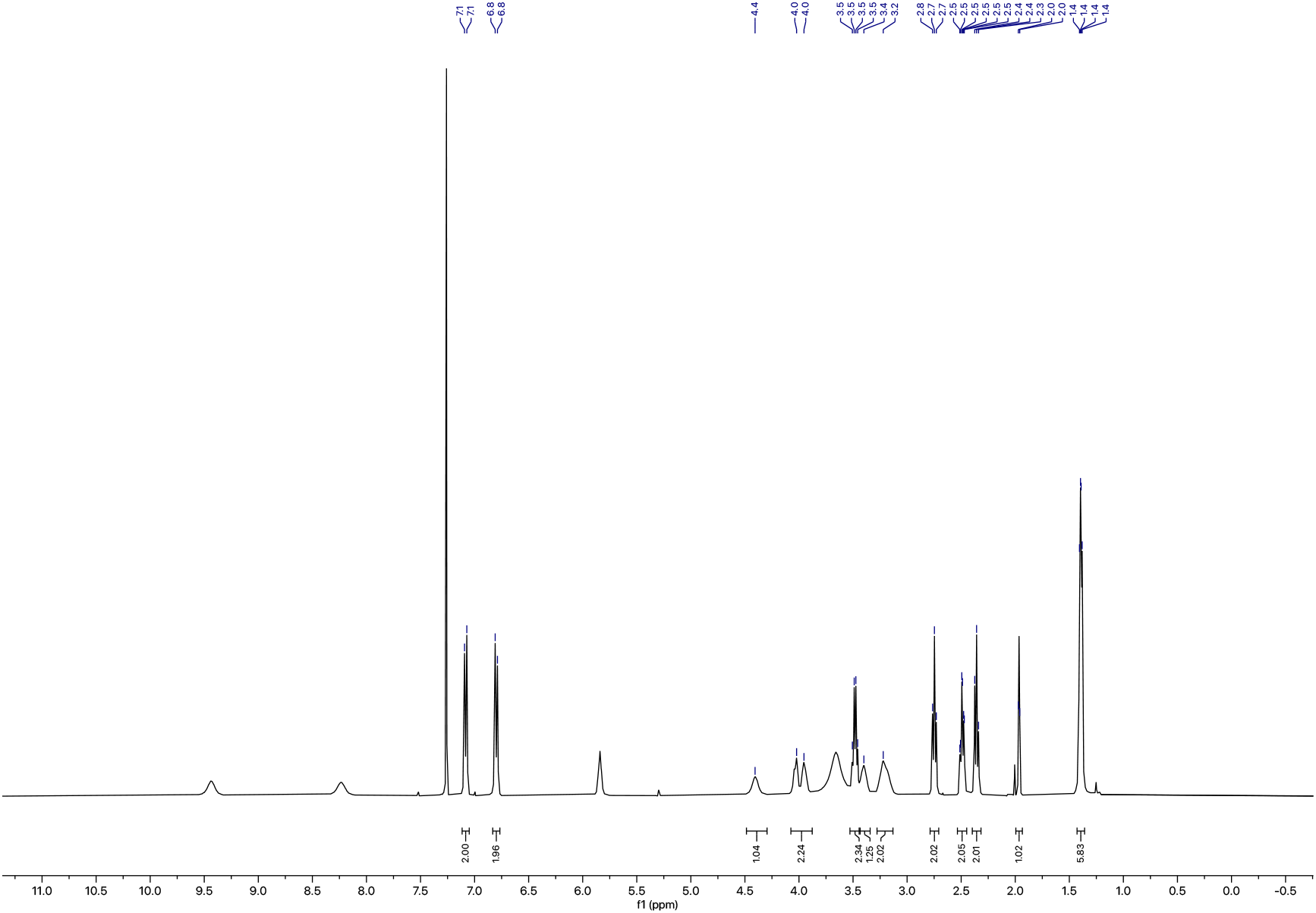

